# Nucleus basalis of Meynert degeneration signals earliest stage of Alzheimer’s disease progression

**DOI:** 10.1101/2023.07.14.547956

**Authors:** Neda Shafiee, Vladimir Fonov, Mahsa Dadar, R. Nathan Spreng, D. Louis Collins

**Author notes:** **Correspondence:** Neda Shafiee, Montreal Neurological Institute, 3801 University Street, Montreal, QC H3A 2B4, Canada.

## Abstract

The Nucleus Basalis of Meynert (NbM) is the main source of cholinergic projection to the entorhinal cortex (EC) and hippocampus (HC), where acetylcholine plays a key role in their function. Both cholinergic cells of NbM and their receptive targets in the EC and HC show sensitivity to neurofibrillary degeneration during the early stages of Alzheimer’s disease. Precise delineation of the NbM region on T1w scans is challenging due to limited spatial resolution and contrast. Using deformation-based morphometry (DBM), we sought to quantify NbM degeneration along the Alzheimer’s disease trajectory and confirm recent studies suggesting that NbM degeneration happens before degeneration in the EC or HC.

MRI scans of 1447 participants from the Alzheimer’s Disease Neuroimaging Initiative (ADNI1, 2 and GO) were up-sampled to 0.5 mm voxel-size in each direction (0.125 mm^3^) before non-linear registration to an ADNI-based unbiased symmetric average MRI template. A subgroup of 677 amyloid-positive participants were chosen for volume time course analysis where the resulting deformation fields were used to compute a high-resolution Jacobian determinant map for each participant as a proxy for local volume change within NbM, EC, and HC masks. Scores of volume change were corrected for age and sex and z-scored using normative data from cognitively healthy amyloid-negative (CN-) participants (n = 219). Two-sample t-tests were then used to compare the baseline differences between NbM, EC, and HC for amyloid-positive cognitively normal controls (CN+), amyloid-positive early mild cognitive impairment (eMCI), and amyloid-positive late mild cognitive impairment (lMCI). Longitudinal linear mixed-effect models were used to compare trajectories of volume change after realigning all participants into a common timeline based on their cognitive decline.

Results showed a significant cross-sectional difference at baseline between CN+ and eMCI, and eMCI and lMCI for both left and right Z-scored NbM volumes. There was no difference in z-scored EC and HC volumes between CN+ and eMCI groups but results from eMCI and lMCI differed significantly in these regions. Longitudinal analysis, with a focus on the early disease stages showed the earliest deviation from the CN- trajectory in the NbM and HC in the subject-specific time realigned data.

Contrary to the notion that Alzheimer’s disease originates in EC, we found that NbM volume changes earlier in the disease trajectory than EC or HC. Converging evidence from cross-sectional and longitudinal models suggest that the NbM may be a focal target of early AD progression, which is often obscured by normal age-related decline.

## Introduction

Alzheimer’s disease (AD) is a neurodegenerative disorder with a dual proteinopathy as key markers of its pathology: accumulation of amyloid-beta in the form of plaques and hyperphosphorylated-tau in the form of neurofibrillary tangles (NFTs). These changes in the brain lead to irreversible loss of neurons and an eventual decline in cognitive and functional abilities as clinical symptoms of the disease manifest^1, 2^.

AD has a complex temporal evolution and the precise sequence of the spread of neurodegeneration across brain regions, especially in the initial stages, remains unclear. Widely accepted models suggest that Alzheimer’s degeneration starts in the entorhinal cortices and then spreads through the temporoparietal cortex^3^. This is supported by a hierarchical system that stages AD according to tau aggregation^4^, where the accumulation of NFT first appears in entorhinal cortices and hippocampus. However, this theory has been challenged by histological^5–8^ and structural imaging studies^9,10^, focusing on the early pathological changes to cholinergic neurons of the basal forebrain. The main source of cholinergic projections to the cerebral cortex is the magnocellular neurons of the nucleus basalis of Meynert (NbM), the largest cluster of cholinergic cells that constitute the basal forebrain. The NbM is located within a continuous band of structures including the amygdala, hippocampus, and entorhinal cortex which are all at high risk of degeneration due to AD pathology. This anatomical positioning of NbM has been speculated to be the main reason for its vulnerability to NFTs^11^. Postmortem studies have also shown high densities of NFTs in NbM in early and presymptomatic stages of the disease^5^. In addition, Fernandez-Cabello *et al.*^12^ suggested that degeneration of the cholinergic projection system is an upstream event of entorhinal and neocortical degeneration, making NbM a possible biomarker early in the course of the disease.

However, the precise delineation of NbM is difficult due to limited spatial resolution and contrast in MR images. The NbM lacks strict boundaries with adjacent cell groups and its small size poses a challenge for defining this region on common 1mm^3^ isotropic T1w scans. To overcome this challenge, we increased the resolution of our MRI scans, before performing a deformation-based morphometry (DBM) analysis. DBM, unlike voxel-based morphometry (VBM), does not depend on an automated segmentation of the MR data into gray matter, white matter, and CSF; instead it can use image contrast directly as an explicit representation of these distributions^13, 14^. The improvements in nonlinear image registration algorithms allow for matching the images locally based on similarities in contrast and intensities, making DBM more sensitive than VBM for subtle differences and more resilient to erroneous tissue classification.

In this study, we precisely quantify volume loss in these structures (NbM, EC, HC) across Alzheimer’s disease trajectory. We look at cognitively normal controls (CN), including those that are amyloid-negative (CN-) and those that are amyloid-positive (CN+), as well as amyloid-positive early mild cognitive impairment (eMCI), amyloid-positive late mild cognitive impairment (lMCI) and amyloid-positive patients with dementia due to clinically probable AD. In addition to examining these groups cross sectionally, we also map all subjects into a common disease timeline to compare longitudinal differences between their continuous volume trajectories. We also verify these results by comparing these trajectories in only the CN+ and eMCI groups to ensure the later disease groups are not driving the early findings.

## Materials and methods

### Dataset

Data used in the preparation of this article were obtained from the Alzheimer’s Disease Neuroimaging Initiative (ADNI) database (adni.loni.usc.edu). The ADNI was launched in 2003 as a public-private partnership, led by Principal Investigator Michael W. Weiner, MD. The primary goal of ADNI has been to test whether serial magnetic resonance imaging (MRI), positron emission tomography (PET), other biological markers, and clinical and neuropsychological assessment can be combined to measure the progression of mild cognitive impairment (MCI) and early Alzheimer’s disease (AD).

All ADNI subjects provided informed consent and the protocol was approved by the institution review board at all sites. In this work, we selected subjects from ADNI for which T1 MRI data was available at the baseline visit. To better focus on AD pathology, and to follow those most likely on the Alzheimer’s trajectory, amyloid positivity was also applied as a key inclusion criterion using both amyloid Positron Emission Tomography (PET) scan and CSF biomarker, with a cut-off of 0.79 SUVR (using composite reference region, corresponding to the ADNI standard whole cerebellum-based florbetapir positivity threshold of 1.11) for PET data^15^ or a CSF pTau/Aß markers of more than 0.028 for positivity^16, 17^. From the 1447 ADNI1, ADNIGO, and ADNI2 subjects available, 677 subjects (with 1982 scans) met these criteria. In addition, we selected the 219 cognitively normal controls that were amyloid negative (CN-) for the z-scoring normalization process described below.

### MRI super-sampling

To address the issue of the very small size of the NbM, we decreased the voxel-size of native MRI scans using the upsampling method introduced by Manjón *et al.*^18^ (See Fig. 1). In MR imaging, the common model assumes that low-resolution voxels can be well modeled as the average of the corresponding high-resolution voxel values plus some acquisition noise. To construct the high-resolution image, the method proposed by Manjón *et al.* enforces a structure-preserving constraint as opposed to imposing an arbitrary smoothness constraint: the down-sampled version of the reconstructed image should be the same as the noise-free low-resolution image for all locations. The super-resolution method is an iterative procedure including (1) a patch-based non-local reconstruction step to perform the upsampling and (2) a mean correction step to ensure that the reconstructed high-resolution image remains consistent with the original low-resolution image. This upsampling method was applied in 3D to all T1w MRI scans prior to other pre-processing steps and results were checked visually for wrong outcomes. The up-sampled scans were then cropped around the medial temporal region to keep the dataset volume within a computationally feasible range.

**Figure 1.**
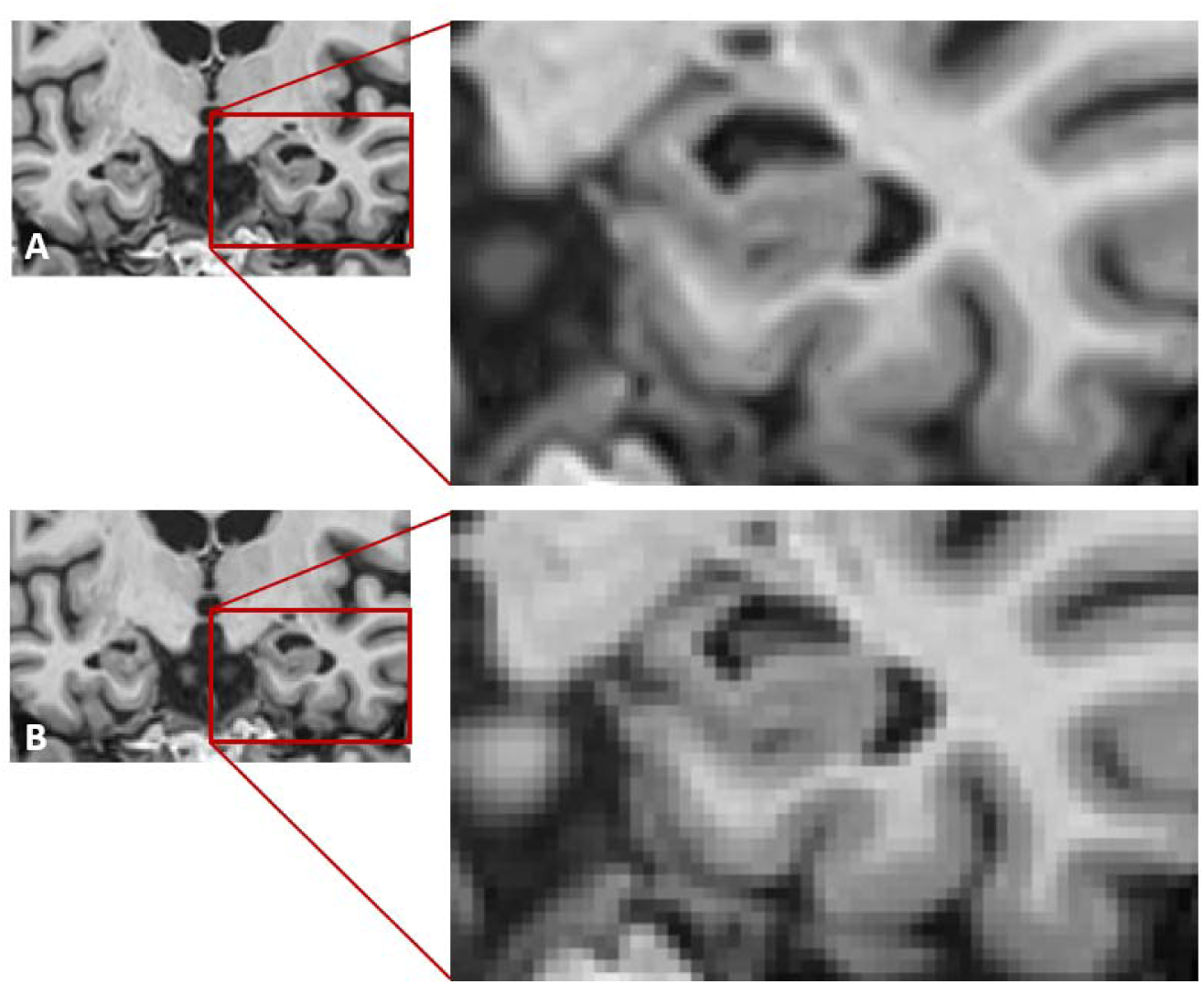
super-resolution results. (A) a super-sampled MRI scan from ADNI dataset and (B) the original low-resolution scan

### MRI Preprocessing

All up-sampled T1w scans were pre-processed through our standard longitudinal pipeline including image denoising^19^, intensity non-uniformity correction^20^, and image intensity normalization into range (0-100) using histogram matching. Each T1w volume was then non-linearly registered to an ADNI-based template using ANTs diffeomorphic registration pipeline^21^. The quality of the registrations was visually assessed and the six cases (1 eMCI, 3 lMCI and 2 AD) that did not pass this quality control were discarded.

### Deformation-Based Morphometry analysis

DBM characterizes positional differences between each voxel of a target image and a standard brain, which then detects morphological differences over the entire brain. Here, each up-sampled subject image was non-linearly registered to an unbiased average template ^22^, resulting in a deformation field as the output. The statistical analyses are then performed on parameters extracted from the deformation fields, instead of the registered voxels. In this study, DBM analysis was performed using MNI MINC tools^23^.

The deformation field D was computed for all subject scans such that the subject S was mapped to the template T when deformed by D (i.e., D(S)). To be able to compare all subjects in the common (template) space, we used the ANTs inverse mapping of D to map the template T to the subject S. The Jacobian matrix of the deformation field is defined as below:

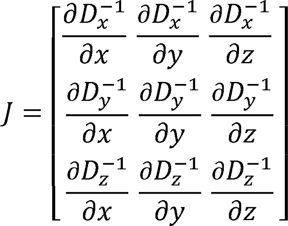

The Jacobian map is then defined as the determinant of the Jacobian Matrix at each voxel. According to this map, areas showing Jacobian determinant values higher than 1, show larger local volume in the subject’s MRI scan relative to the template, while areas of the map with a value less than 1 indicate a smaller local volume.

#### 1.1.1. Atlas-based volumetry

All Jacobian maps were calculated in the template space, thus normalizing for head size. From the atlas published by Zaborszky *et al.* ^24^, we combined both the Ch4 and Ch4p probabilistic areas to define the NbM region. A threshold of 50% was used define the NbM mask ^25^. For HC and EC regions, atlases were created by running ASHS^26^, the software for automatic segmentation of medial temporal lobe regions on the average template volume. Again, mean values of z-scored Jacobian maps in the left and right HC and EC regions for each subject were used (Fig. 2). The mean values of Jacobian maps inside the left and right masks were then calculated for each subject.

**Figure 2.**
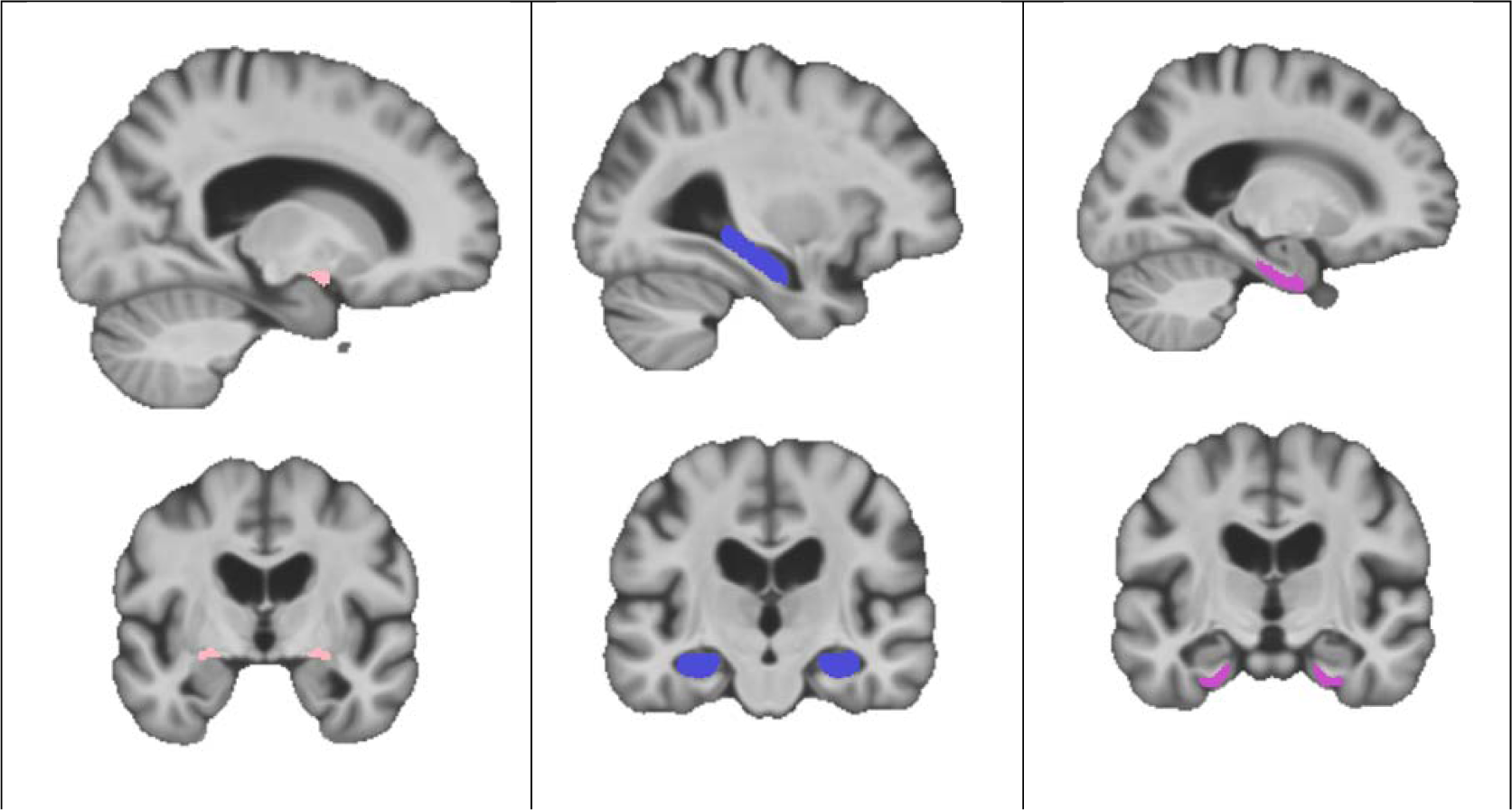
regions of interest. NbM (left), Hippocampus (middle), and Entorhinal cortex (right).

### Longitudinal analysis

#### Subject-specific template

To better assess the trajectory of anatomical changes over the course of the disease, we also incorporated follow-up scans into our analysis. Longitudinal image processing aims at reducing within-subject variability by transferring information across time, e.g., enforcing temporal smoothness or informing the processing of later time points with results from earlier scans. However, these approaches are susceptible to processing bias that could be due to non-linear registration when using an arbitrary reference image such as a general template. Consistently treating a single time point, usually baseline, differently from others, for instance, to construct an atlas registration or to transfer label maps for initialization purposes, can already be sufficient to introduce bias ^27^.

It is unlikely that bias affects all groups equally, considering that one group usually shows only little longitudinal change, while the other undergoes significant neurodegeneration. On the other hand, using an individual template per subject has been shown to effectively eliminate bias ^27^. The approach which is based on the work done by Fonov *et al.* ^28^ treats all data-points equally without prioritizing baseline data over the follow-up scans and has been utilized to obtain a more accurate anatomical correspondence between time-points.

To summarize, the objective of the subject-specific template creation algorithm is to find the non-linear transformations that minimize the anatomical shape differences between all-time points to create the most representative average of the subject’s anatomy, where we expect a 1:1 anatomical correspondence throughout the brain. Processing is achieved in two steps. First, all data is processed cross-sectionally to bring each volume into stereotaxic space. Second, this data is used to build a subject-specific individual template.

After building the subject-specific template, we calculate the local volume change in each data-point compared to the subject-specific template, resulting in an unbiased relative volume change based on time. To make sure that longitudinal change for each subject is comparable with the other subjects, the measured change is scaled using the difference between each subject-specific template and a general ADNI template (Figure 3). The ADNI template used for this work is made up of 150 ADNI subjects (50 cognitively normal, 50 MCI, and 50 dementia due to AD). The results were then corrected for age and sex using the healthy amyloid-negative subjects as the control group.

**Figure 3.**
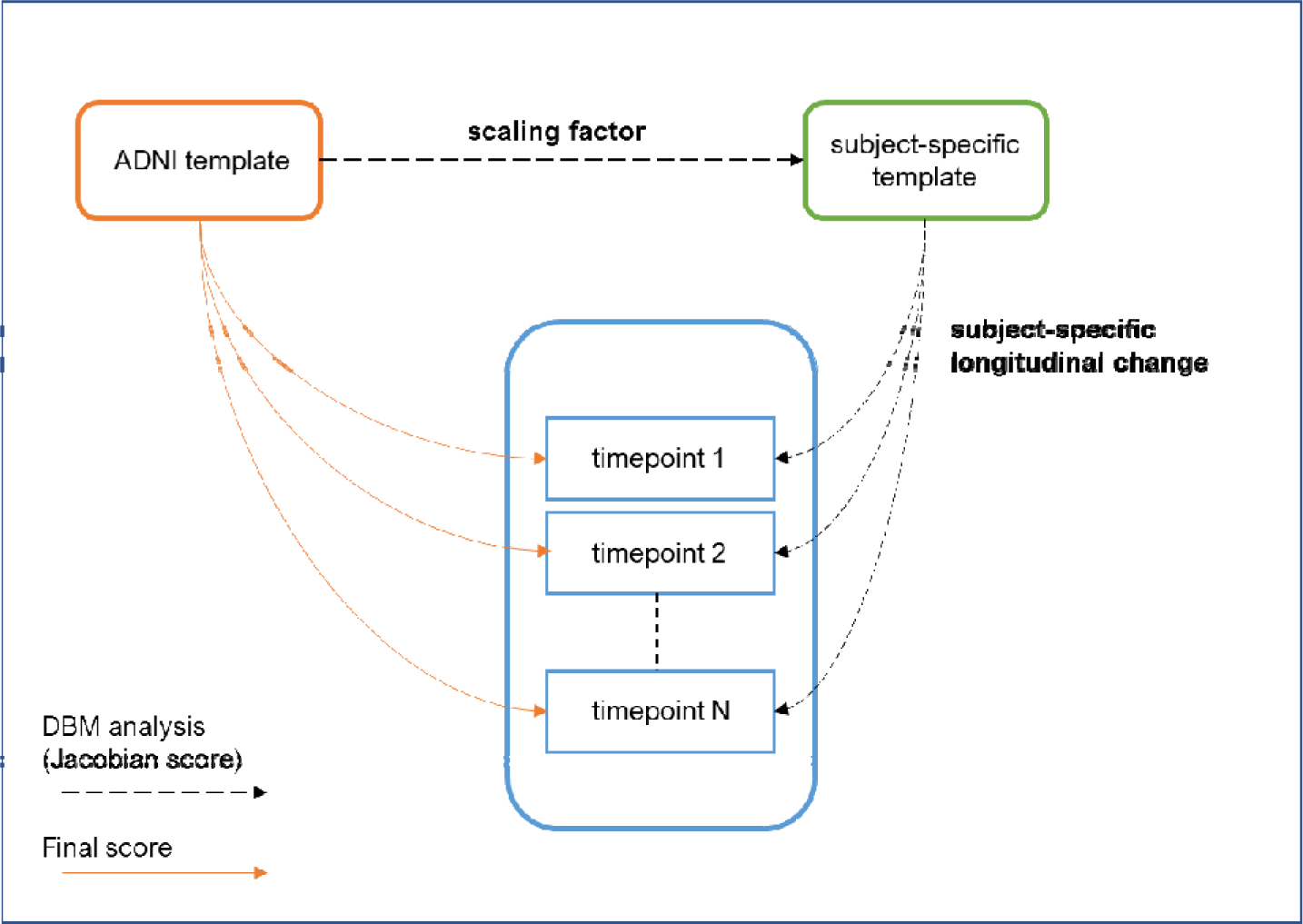
Simplified diagram of the method used to calculate overall volume change in reference to an ADNI template.

#### Subject-specific time-shift

When patients get their diagnosis, whether it is early or late MCI or clinical AD, it is impossible to determine exactly when the disease started. Moreover, not all subjects progress in the same manner and rate. However, clinical measurements such as cognitive scores can - to some extent - estimate where in the disease trajectory each patient stands, with a time scale dependent on the clinical or cognitive measures used. To address the issue of estimating the time of disease onset, we investigated a new technique that leverages cognitive test scores to estimate a latent timeline that models the subject-specific disease progression.

Here we used the work of Kühnel *et al.*^29^ to model the progression of Alzheimer’s disease using cognitive scores at different time points. In this work, a nonlinear mixed-effects model aligns patients based on their predicted disease progression along a continuous latent disease timeline. More specifically, the Alzheimer’s Disease Assessment Scale-cognitive subscale (ADAS-cog-13) and the Mini-Mental State Examination (MMSE) were used simultaneously to fit an exponential curve to the longitudinal data, calculating time-shifts for each subject. Unlike the work in Kühnel *et al.*, where the diagnostic group of each subject was added to the mixed-effect model resulting in a group-based difference for the latent timeline, we did not include this label in our analysis, making the model blind to any previous classification and hence, purely data driven.

Figure 4 depicts the results of MMSE and ADAS13 scores plotted against the original follow-up time (left column), and the subject-level estimated time-shifts (right column). Longitudinal data from 219 cognitively normal amyloid negative subjects (CN-) was included with the 677 amyloid positive subjects before running the analyses to ensure a common time scale across all subjects for later analysis.

**Figure 4.**
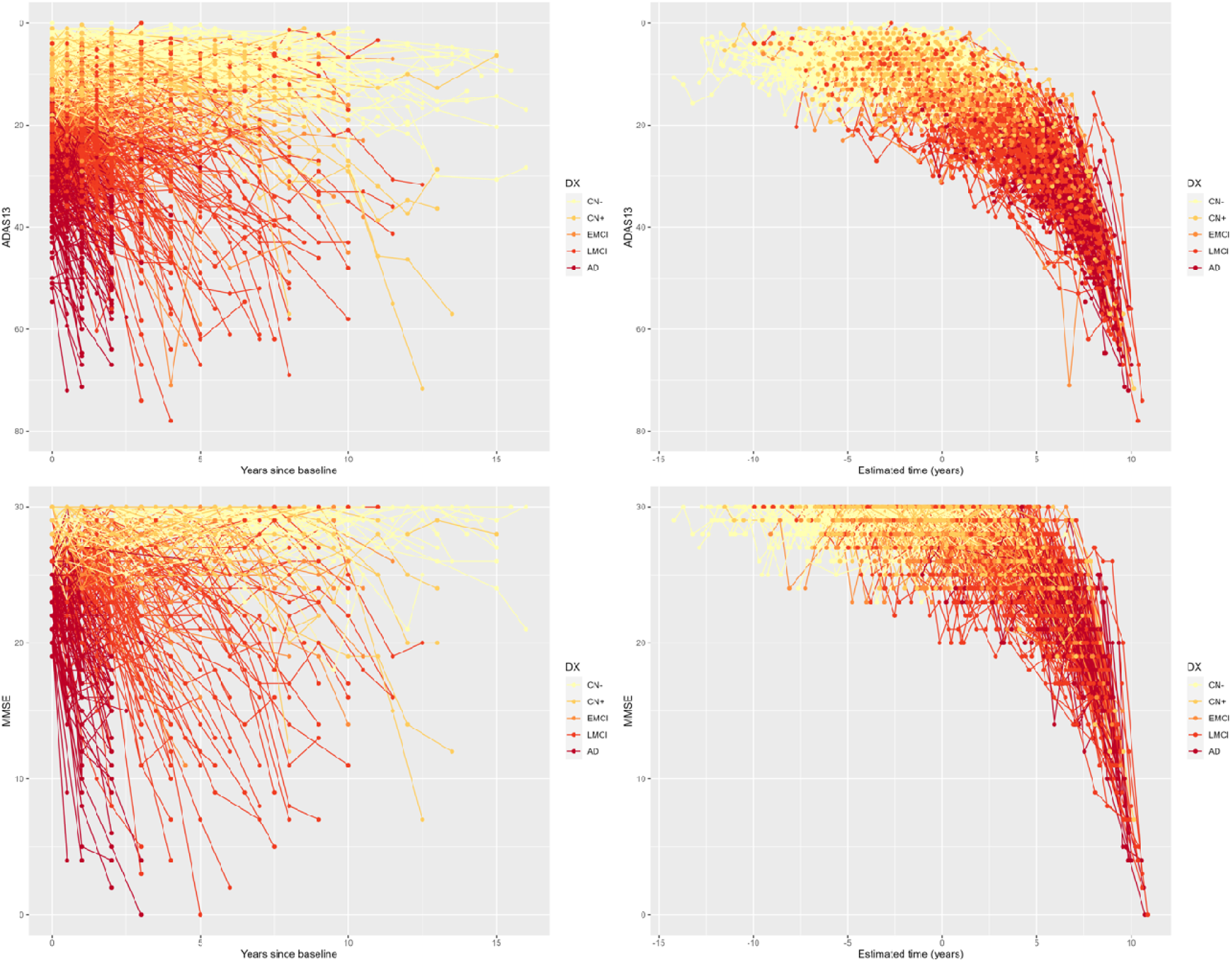
Predicted patient staging using MMSE and ADAS-Cog-13 scores. The horizontal axis is the time scale predicted by the model that is adjusted for subject-level differences in disease stage, thus bringing all subjects onto a common timeline.

Additionally, and for the sake of comparison, we repeated the exact same steps as Kühnel’s ^29^ and included the group-based difference and found that our results were comparable. Assuming the group-based time-shift for cognitively normal subjects is zero, the average time-shift based on MMSE and ADAS13 for the eMCI group was 36.8 months (about 3 years) from a latent time origin. For lMCI, this shift was 87.5 months (about 7 and a half years) and patients with clinically diagnosed AD had 128.8 months (about 10 and a half years) of shift in time.

#### Statistical analysis

With both subject-specific estimated time and longitudinal local change in the volume, we plotted the trajectories of three regions of interest (HC, EC, and NbM) across the disease progression timeline. Longitudinal modeling of atrophy was performed using a linear mixed effect model to account for both within-subject variability of volumes over multiple scans and between subject changes. All statistical analyses were done using the lme4 package^30^.

The estimated disease time (EDT) was used as the fixed effect (including an intercept and both a linear and quadratic compound). A cubic time effect was also tested but did not improve the model. After evaluation with an Akaike criterion, the final model used the following formulas:

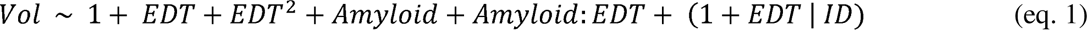

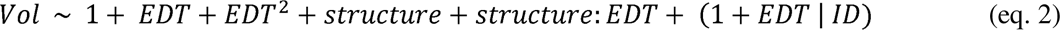

In equation (1), *Amyloid* is a categorical variable showing amyloid positivity vs. negativity in participants. This would compare the time-based trajectory of all amyloid positive participants with those of control normal group (CN-). The term *structure* in the second equation shows the ROI under study (NbM, EC or HC). This equation is added to compare the curves of NbM with those of HC and EC only for amyloid positive participants.

In a second step, to study the effect of APOE status and sex on the trajectories, these variables were added to the model as fixed effects. The model was then calculated as follows:

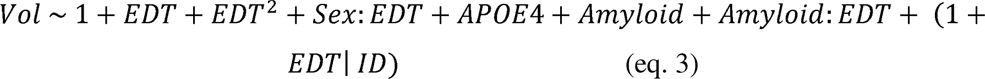

The intercept and slope for each subject were considered as random effects. The term *Amyloid* is a categorical variable showing amyloid positivity vs. negativity (same as eq. 1).

## Results

### Demographics

Table 1 shows the demographics of the 5 groups included in this study. There was a significant age difference between amyloid-positive normal controls and subjects with early-stage mild cognitive impairment (*p*-value<0.001, *t-*value=4.25, *df*=246). The age difference was not significant for other groups.

**Table 1.**
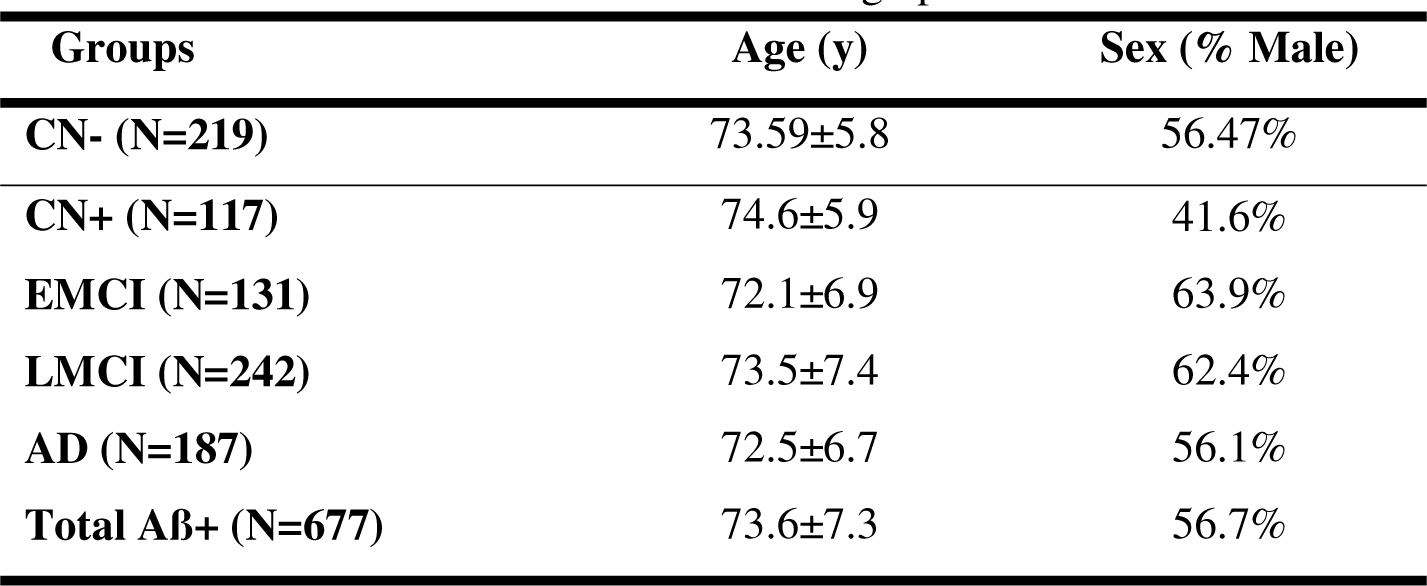
Baseline demographic

| Groups | Age (y) | Sex (% Male) |
| --- | --- | --- |
| CN- (N=219) | 73.59±5.8 | 56.47% |
| CN+ (N=117) | 74.6±5.9 | 41.6% |
| EMCI (N=131) | 72.1±6.9 | 63.9% |
| LMCI (N=242) | 73.5±7.4 | 62.4% |
| AD (N=187) | 72.5±6.7 | 56.1% |
| Total AB+ (N=677) | 73.6±7.3 | 56.7% |

### Atlas-based DBM analysis of baseline data

Figure 5 shows the distribution of z-scored values of mean DBM Jacobian at baseline for NbM, EC, and HC regions, for CN-, CN+, eMCI, lMCI, and patients with clinically probable AD. After Bonferroni correction for multiple comparisons, a significant difference was found between CN+ and eMCI groups for left, and more substantial on right NbM. Additionally, a significant difference was observed between eMCI and lMCI groups, where the difference was more substantial in the left NbM.

**Figure 5.**
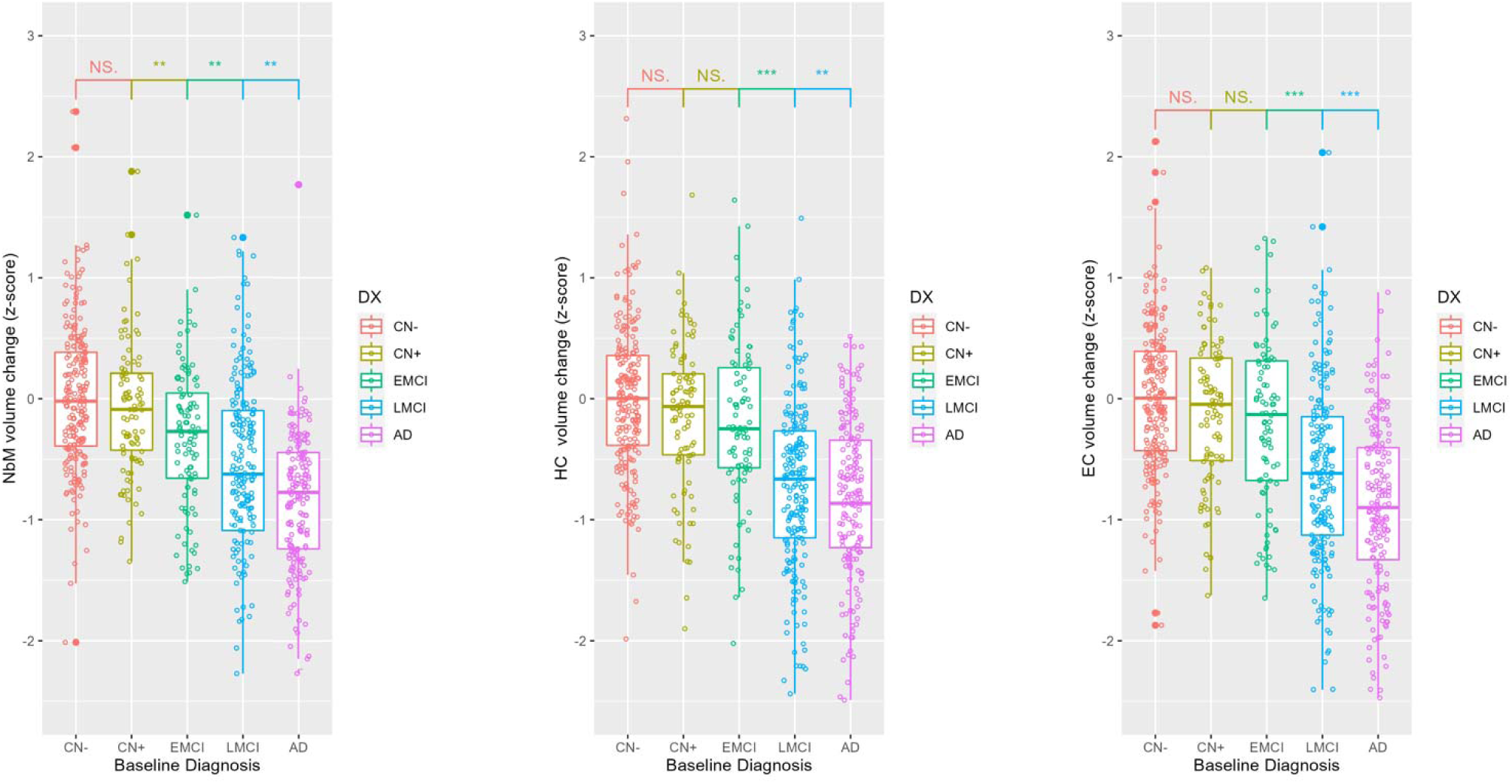
Mean z-scored Jacobian value. Scores for average NbM (left), HC (middle) and EC (right) (***: p-value<5e-06; **: p-value<0.005; *: p-value<0.05; NS: Not significant)

The z-scored EC and HC volumes did not show a statistically significant difference between CN+ and EMCI groups; However, results from eMCI and lMCI differ significantly in these regions for both the left and right hemispheres. All p-values are reported in Table 2. We also observed statistically significant differences between lMCI and AD in NbM, EC, and HC (*P*<0.001). The results are also depicted in Fig. 5 for all participants, where left and right ROIs are combined to obtain average bilateral measurements.

**Table 2.** Statistical differences of atlas based DBM scores between diagnostic groups

| Region | CN+ vs eMCI |  |  | eMCI vs IMCI |  |  |
| --- | --- | --- | --- | --- | --- | --- |
|  | Cohen's *d* | *t*-value | *p*-value | Cohen's *d* | *t*-value | *p*-value |
| Right NbM | 0.38 | 3.087* | 0.0089 | 0.4 | 3.601** | 0.00038 |
| Left NbM | 0.33 | 2.247* | 0.0258 | 0.47 | 4.447** | 1.261e-05 |
| Right EC | 0.06 | 0.4778 | 0.6332 | 0.68 | 6.0101*** | 7.002e-09 |
| Left EC | 0.05 | 0.3878 | 0.6985 | 0.71 | 6.3251*** | 1.255e-09 |
| Right HC | 0.15 | 1.167 | 0.2446 | 0.73 | 6.266*** | 1.953e-09 |
| Left HC | 0.12 | 0.916 | 0.3605 | 0.81 | 7.067*** | 1.906e-11 |

### Longitudinal analysis

The longitudinal data used in this work included 1982 scans from the 677 amyloid-positive and 219 cognitively normal amyloid-negative subjects. On average, each subject had ∼3 time-points. For the CN+ group, the average time-points available per subject was 3.27, for eMCI this average was 2.8, for lMCI, 3.68 and for subjects with AD dementia, the average was 2.3 scans. Using the latent time shift (Fig. 4) to offset the baseline (and following) scans, we were able to plot the changes in three different regions of the brain (NbM, EC and HC) across the estimated disease progression timeline (see Fig. 6 (bottom row) and 7). Note that for longitudinal analysis, left and right ROIs are combined to obtain bilateral measurements. Figure 6 shows how the estimated time-shift, changes the shape of the plots whenever it was used instead of age as a continuous measure of time.

**Figure 6.**
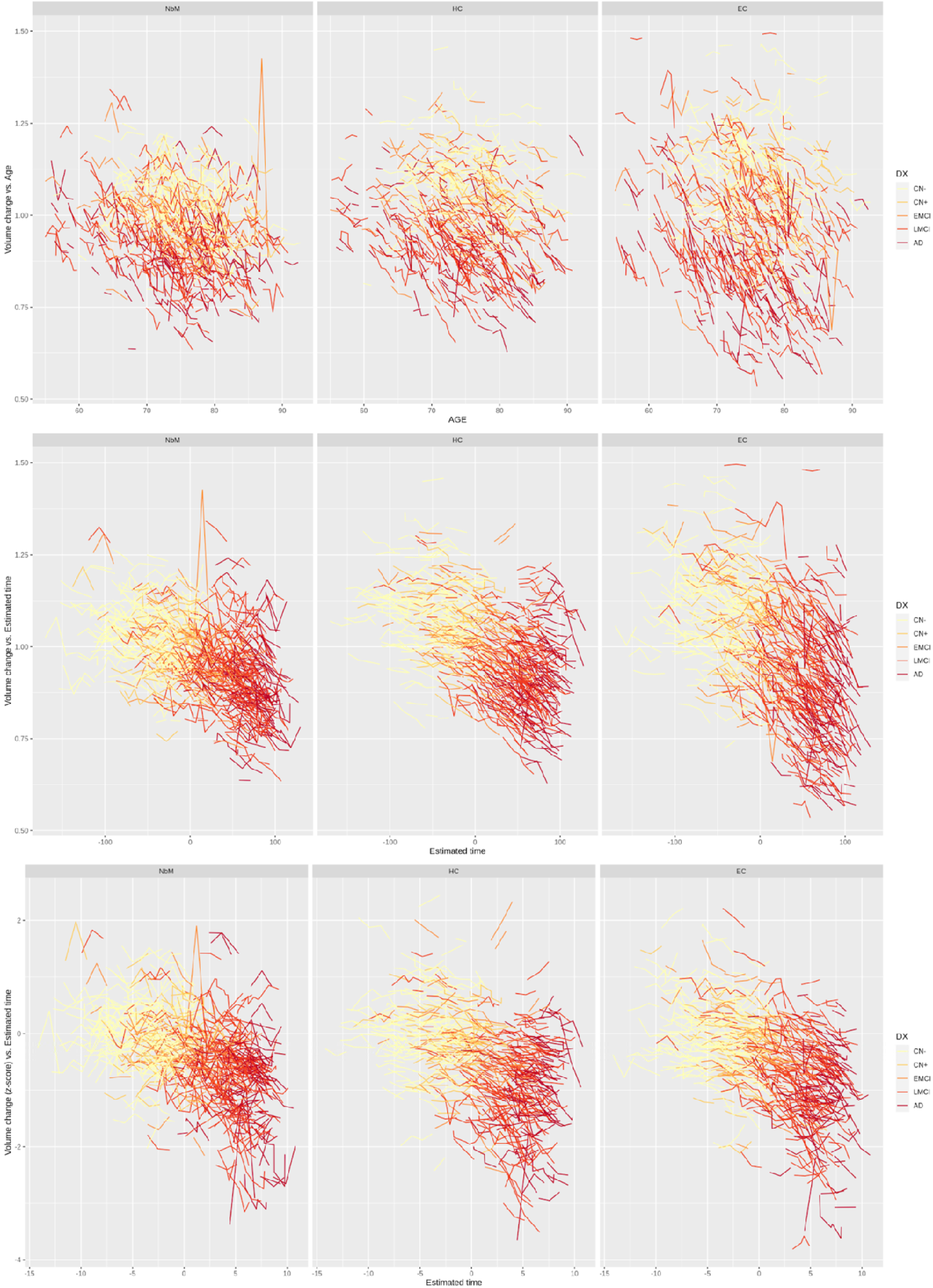
Volume change in NbM, HC, and EC. Volume changes across the age (top row), across the estimated disease timeline (middle row) and the across the estimated disease timeline after z-score normalizing the volume measurements for age and sex (bottom row).

The results of the statistical analyses are also provided in Table 3, where the fitted beta values and statistical significance for eq. 1 is reported.

**Table 3.** Results for statistical analyses (eq. 1)

| Fixed effects | Nucleus basalis |  | Hippocampus |  | Entorhinal cortex |  |
| --- | --- | --- | --- | --- | --- | --- |
|  | Estimate | *p*-value | Estimate | *p*-value | Estimate | *p*-value |
| Intercept | -0.1010 | 0.0494* | -0.5982 | 0.184767 | -0.1446 | 0.000828 *** |
| EDT | -0.3187 | 5.19e-06 *** | -0.1555 | 0.010896 * | -0.3739 | 1.62e-11 *** |
| EDT<sup>2</sup> | -0.2365 | 4.24e-08 *** | -0.1001 | 0.005968 ** | -0.1873 | 2.19e-08 *** |
| Amyloid (positive) | -0.4479 | 1.42e-13 *** | -0.3599 | 1.25e-11 *** | -0.2299 | 5.50e-06 *** |
| Amyloid:EDT | -0.3326 | 0.000107 *** | -0.2569 | 0.000742 *** | -0.1876 | 0.006084 ** |

As can be seen from both Figure 7 and Table 3, when looking into all longitudinal data, EC shows a steeper decline across the trajectory, followed by NbM, and then HC (this can be calculated using the first derivative of the eq. 1 at EDT=0 (early-stage), where the slope equals the coefficient of the EDT in the equation). However, our mixed model shows a more pronounced difference between the cognitively normal amyloid-negative group and combined amyloid-positive groups in NbM compared to EC (and HC). In addition, the Amyloid:EDT term shows that volume loss over time related to amyloid positivity is greater - both numerically and statistically - in the NbM compared to the HC and EC. Furthermore, cognitively normal amyloid-negative group and combined amyloid-positive groups seem to start differentiating earliest in the HC, closely followed by the NbM and then the EC.

**Figure 7.**
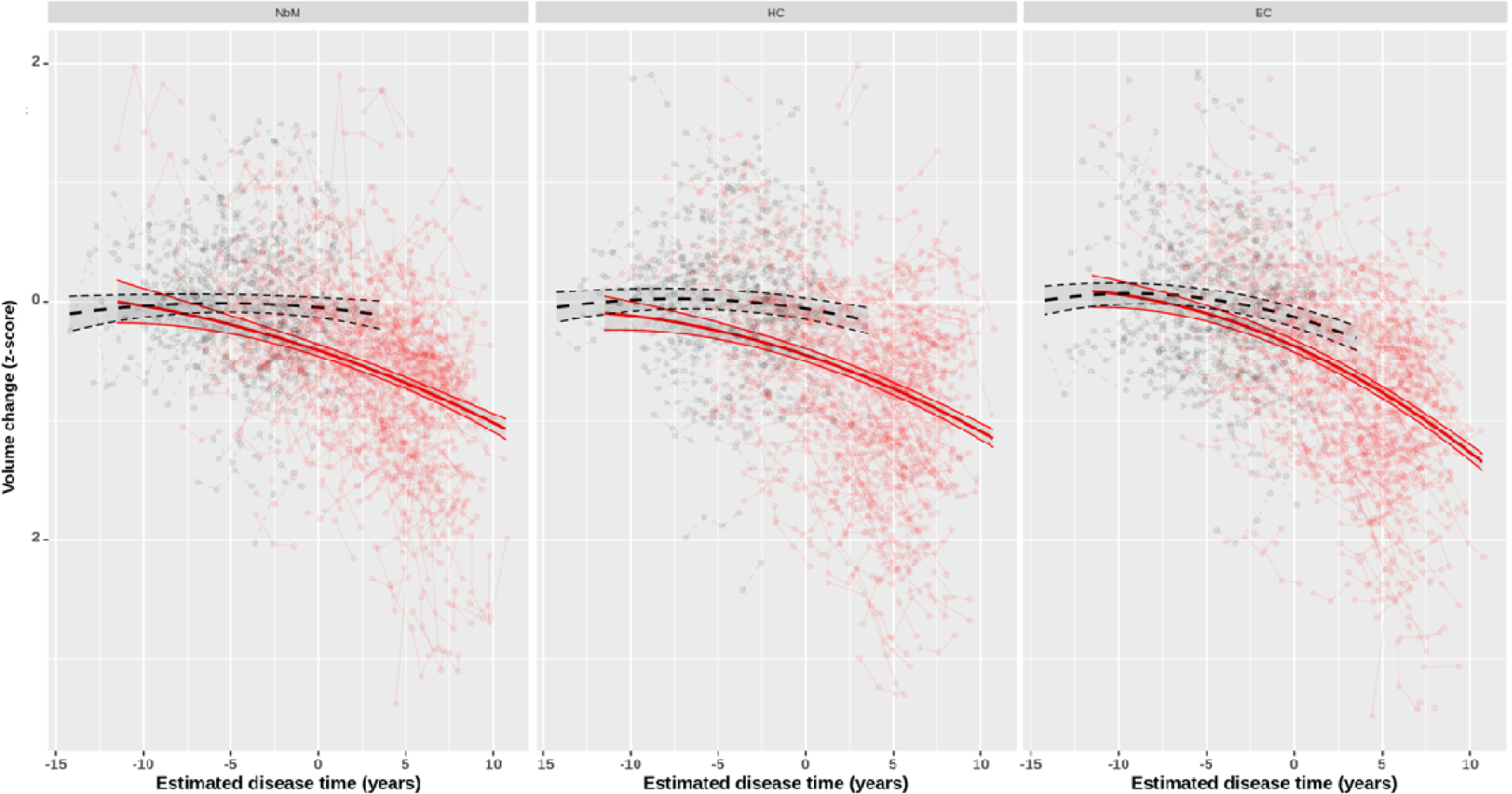
Longitudinal age- and sex-corrected zscore volumes for NbM, HC, and EC. (left to right) across the predicted timeline (estimated using eq. 1). Black lines indicate the CN- group and red shows all amyloid-positive subjects from CN+, eMCI, lMCI and AD subjects.

### Early-stage longitudinal analyses

To ensure that the fitted curves in Figure 7 are not driven by the late-stage atrophy of the lMCI and AD group, we opted to focus on the only the CN+ and eMCI ‘early-stage’ subjects, comparing their trajectories to the CN- group. Figure 8 shows the age- and sex- corrected z-scores for both CN- and the *early-stage* CN+ and eMCI groups for NbM, EC and HC. This time results show a steeper and earlier decline in the *early-stage* CN+ subjects in NbM atrophy trajectory compared to HC and EC. Additionally, no statistically significant differences were observed between normal and abnormal subjects for EC and HC. Age- and sex-normalized NbM and HC trajectories for CN- remains flat, while the CN- trajectory for EC declines with time. This results in greater volume differences for NbM and HC between the CN- group and the CN+ and eMCI groups across the disease time spectrum estimated.

**Figure 8.**
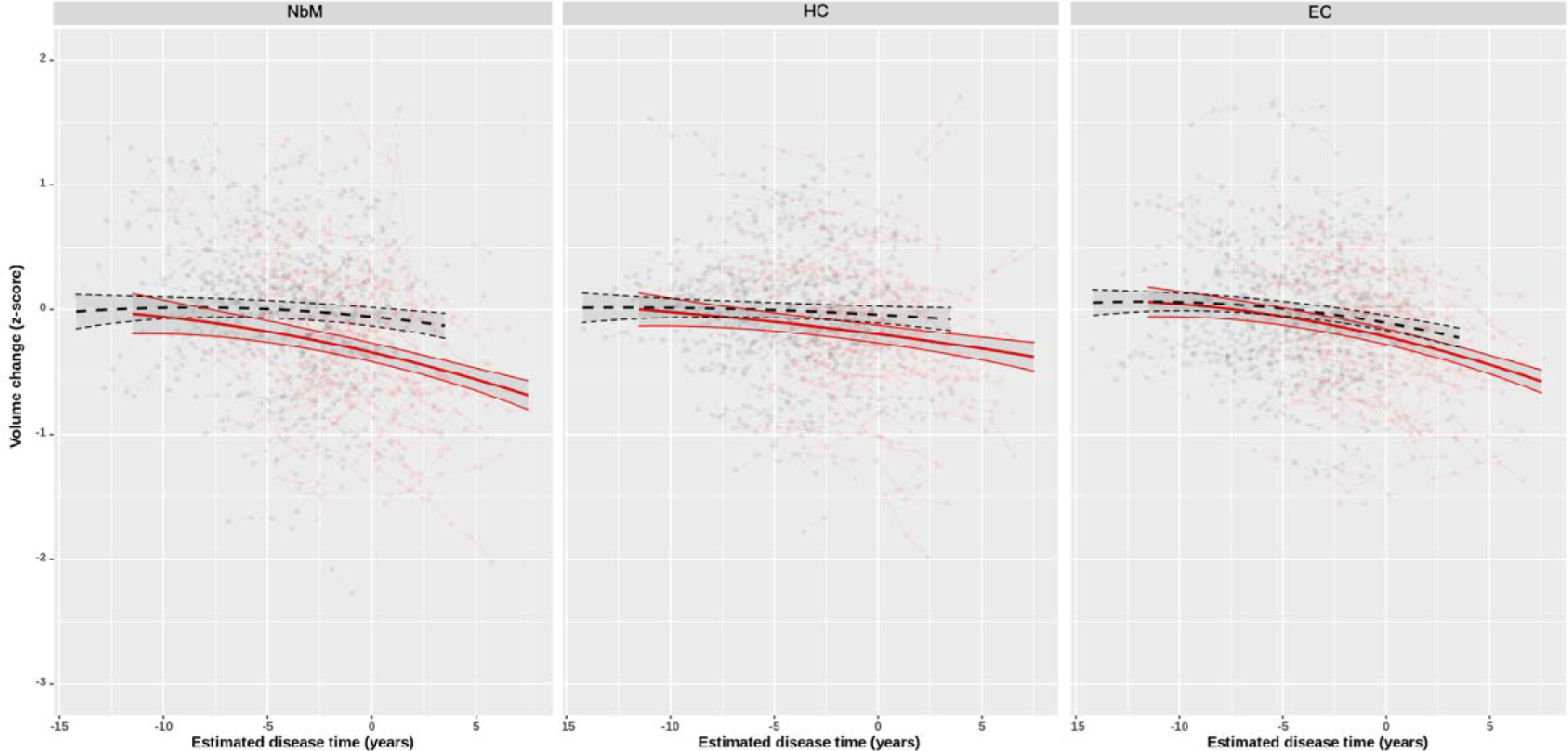
Early-stage volume change. Longitudinal volume change in NbM, HC, and EC (left to right) across the predicted disease timeline (estimated using eq. 1). For each structure for CN- (black) and the early-stage CN+ and eMCI groups (red).

The results of the statistical analyses for the early-stage analysis are also provided in tables 4 and 5. Table 4 shows the fitted beta values and statistical significance for eq. 2. This time instead of comparing the trajectories of normal and abnormal subjects, we opted to compare the abnormal curves of NbM with those of HC and EC. using eq. 2. Results show the effect of EDT, anatomical *structure* and the interaction term (EDT:*structure*) on the longitudinal data. The fitted model shows that the normalized volume difference at EDT=0 between the NbM and the HC is significant (*p*=0.0166), and between the NbM and the EC is almost significant (*p*=.055). The slope of the trajectory between NbM is statistically steeper than both the EC (*p*=.015) and the HC (*p*<.001).

**Table 4.** Results for early-stage analyses (eq. 2)

| Fixed effects | Estimate | *p*-value | *t*-value |
| --- | --- | --- | --- |
| Intercept | **-0.1297** | **2.56e-07 \*\*\*** | **-5.933** |
| EDT | **-0.5304** | **1.48e-11 \*\*\*** | **-7.075** |
| EDT<sup>2</sup> | -0.2273 | 0.16206 | -1.399 |
| EC vol (vs. NbM) | 0.031 | 0.05495 . | 1.919 |
| HC vol (vs. NbM) | **0.051** | **0.00166 \*\*** | **3.147** |
| EDT:EC vol | **0.17737** | **0.01529 \*** | **2.425** |
| EDT:HC vol | **0.37425** | **3.25e-07 \*\*\*** | **5.121** |

**Table 5.** Results for early-stage analyses (eq. 3)

| **Fixed effects** | **Nucleus basalis** |  | **Entorhinal cortex** |  | **Hippocampus** |  |
| --- | --- | --- | --- | --- | --- | --- |
|  | Estimate | *p*-value | Estimate | *p*-value | Estimate | *p*-value |
| Intercept | -0.0872 | 0.0623. | **-0.1401** | **0.00295 \*\*** | -0.0552 | 0.1486 |
| EDT | **-0.3298** | **0.0005\*\*\*** | **-0.3408** | **2.27e-08 \*\*\*** | **-0.2406** | **0.0146 \*** |
| EDT<sup>2</sup> | **-0.3431** | **0.0014\*\*** | **-0.1987** | **0.0164 \*** | -0.1390 | 0.2615 |
| Amyloid (positive) | **-0.4260** | **6.37e-08 \*\*\*** | **-0.2122** | **0.0012 \*\*** | **-0.208** | **0.0025 \*\*** |
| Amyloid:EDT | **-0.2938** | **0.0011 \*\*** | **-0.1834** | **0.0074 \*\*** | -0.1527 | 0.05709. |
| Sex (male):EDT | 0.0568 | 0.4015 | **0.1598** | **0.00244 \*\*** | **0.1616** | **0.013 \*** |
| APOE4 status (1) | -0.1427 | 0.0574. | -0.0339 | 0.53074 | -0.0620 | 0.3583 |
| APOE4 status (2) | -0.1213 | 0.0505. | -0.1344 | 0.41141 | -0.1218 | 0.3448 |

Table 5 shows the effect of EDT, amyloid status, sex:EDT and APOE4 status on the volume trajectories of NbM, EC and HC when looking at early-stage dataset using eq. 3. We see that EDT is significant for all three structures, but that EDT^2^ is significant only for NbM and EC. Amyloid positivity results in smaller volumes for all three structures, and over time, amyloid positivity results in a significantly steeper decline for NbM (*p*=.0011, *t*-value = 3.149) and EC (*p*=.0074, *t*-value = −2.752) but is only trending for HC (*p*=.057, *t*-value = 1.904). There was a trend for greater atrophy of the NbM in subjects with the APOE-4 allele.

## Discussion

In this work, we were able to quantify the volume changes in NbM, EC and HC regions due to Alzheimer’s disease on T1w scans. Our cross-sectional results show that while macroscopic changes in EC and HC emerge as MCI subjects advance toward the later stages, NbM degeneration starts sooner, with atrophy apparent in the preliminary stages of early MCI. This points to MRI-based measurements of NbM as potential biomarkers for early detection of AD and as a marker of disease burden. The findings of the cross-sectional analysis are in line with previous studies working on NbM and Basal Forebrain^12^. It has been long-established that NbM, the source of cholinergic innervation, undergoes severe neurodegeneration in Alzheimer’s disease. In more recent studies, Schmitz *et al.*^31^ have provided evidence that basal forebrain pathology precedes and predicts both entorhinal pathology and memory impairment. Our findings regarding AD-related atrophy in both EC and HC were consistent with previous studies, confirming the neurodegeneration in these regions during the mild cognitive impairment stage.

The outcome of longitudinal analysis confirms the cross-sectional finding that NbM undergoes neurodegeneration earlier than regions such as EC and HC. Once subjects are realigned in time with a latent offset starting point for the disease, we find that the NbM is the earliest structure to be affected by neurodegeneration measured by atrophy and is the first structure to be associated with early cognitive changes, even though the disease time offset was driven by ADAS13 and MMSE, tests that are linked more to memory than to attention. It is important to consider the very large heterogeneity of disease burden and cognitive decline in the Alzheimer’s trajectory.

In this work we used a new method of sequencing the disease stage in a continuous manner, using the clinical symptoms instead of using age as the time-based variable. Our results of subject-specific disease time analysis are comparable with the original work of Kühnel^29^ on subjects from the ADNI dataset. In their work, they found ∼25 months of time-shift for subjects with memory concern (we did not include this group), ∼50 months for eMCI (36.8 months in our work), ∼90 months for lMCI (87.5 months in our work) and ∼150 months for subjects with dementia (128.8 months in our work). The differences may be explained by Kühnel’s use of the CDR score while we used ADAS-Cog13 and MMSE to drive the latent time offset as they are more sensitive to subtle cognitive changes. This new method enabled us to follow the trajectory of the disease progression in macroscopic MRI-based measurements, independent of age (as an indirect measure of disease progression) or diagnosis group (which provides a discrete timeline).

The result of our work is in line with previous studies on the association of hippocampal and entorhinal atrophy with sex differences. Two studies on ADNI and MIRIAD determined that atrophy rates were faster by 1–1.5% per year in women with aMCI and AD dementia than in men ^32, 33^. However, while we found a sex:EDT effect for HC and EC over and above that expected for normal age and sex, we were unable to see the effect of sex on NbM atrophy rates.

To ensure that more pronounced volume changes in the later-stages of the disease were not driving the measurements for earlier stages through an artefact of model fitting, we defined a new group consisting only of cognitively normal amyloid positive subjects and those with early MCI, referred to as “early-stage”. We then repeated our statistical analyses on this group and compared the longitudinal changes with healthy amyloid negative subjects. The results showed that even in the early stage of AD, NbM shows a steeper atrophy trajectory compared to HC and EC, and this difference becomes more pronounced as the disease progresses.

This study is not without limitations, and this should be considered when interpreting our findings. First, our use of Jacobian maps as an indirect measure of atrophy may be susceptible to misregistration or measurement errors despite our pipeline’s robustness. However, all nonlinear registrations were visually assessed and cases that did not pass this quality control step were removed. Additionally, although we used the most widely used atlas, defining the neuroanatomical boundaries of the nucleus basalis of Meynert is inherently difficult, given the cluster-like nature of this anatomical region. This difficulty is also reflected in the limited spatial consistency of the NbM between different published atlases^25^. Another important point to consider is the sex imbalance in the different subject groups used here. As shown previously in the data section, data from female subjects drive the measurements in the CN+ group, and for MCI groups, male subjects are in the majority. Moreover, we’re not taking into consideration that there might be different patterns of atrophy for subtypes of patients, especially when looking into the whole timeline, but early-stage might be too soon to see the differences between patterns, keeping our results robust. Finally, we used cognitive scores to estimate the time of the disease; however, damage to the NbM is primarily associated with impairment of attention, which is not well assessed in ADNI. Thus, although the time realignment of the subjects is roughly correlated with the disease start point, such time offsets might underestimate the onset of atrophy in the NbM. Nonetheless, using measures of global cognition instead of focusing on attention ensures a more unbiased approach for comparing these three structures.

The results of this study provide additional support for a subcortical onset of AD. Accurate disease staging, prior to the emergence of a memory impairment, is critical for the development of biomarkers and novel intervention targets.

## Acknowledgements

Data collection and sharing for this project was funded by the Alzheimer’s Disease Neuroimaging Initiative (ADNI) (National Institutes of Health Grant U01 AG024904) and DOD ADNI (Department of Defense award number W81XWH-12-2-0012). ADNI is funded by the National Institute on Aging, the National Institute of Biomedical Imaging and Bioengineering, and through generous contributions from the following: AbbVie, Alzheimer’s Association; Alzheimer’s Drug Discovery Foundation; Araclon Biotech; BioClinica, Inc.; Biogen; Bristol-Myers Squibb Company; CereSpir, Inc.; Cogstate; Eisai Inc.; Elan Pharmaceuticals, Inc.; Eli Lilly and Company; EuroImmun; F. Hoffmann-La Roche Ltd and its affiliated company Genentech, Inc.; Fujirebio; GE Healthcare; IXICO Ltd.; Janssen Alzheimer Immunotherapy Research & Development, LLC.; Johnson & Johnson Pharmaceutical Research & Development LLC.; Lumosity; Lundbeck; Merck & Co., Inc.; Meso Scale Diagnostics, LLC.; NeuroRx Research; Neurotrack Technologies; Novartis Pharmaceuticals Corporation; Pfizer Inc.; Piramal Imaging; Servier; Takeda Pharmaceutical Company; and Transition Therapeutics. The Canadian Institutes of Health Research is providing funds to support ADNI clinical sites in Canada. Private sector contributions are facilitated by the Foundation for the National Institutes of Health (https://fnih.org/). The grantee organization is the Northern California Institute for Research and Education, and the study is coordinated by the Alzheimer’s Therapeutic Research Institute at the University of Southern California. ADNI data are disseminated by the Laboratory for Neuro Imaging at the University of Southern California.

